# Nerve fibers in the Tumor Microenvironment are co-localized with Tertiary Lymphoid Structures

**DOI:** 10.1101/2020.08.07.232322

**Authors:** Lara R. Heij, Xiuxiang Tan, Jakob N. Kather, Jan M. Niehues, Shivan Sivakumar, Nicole Heussen, Gregory van der Kroft, Steven W.M. Olde Damink, Sven Lang, Jan Bednarsch, Merel R. Aberle, Tom Luedde, Nadine T. Gaisa, Drolaiz H.W. Liu, Jack P.M. Cleutjens, Dominik P. Modest, Georg J. Wiltberger, Ulf P. Neumann

**Affiliations:** Department of General, Gastrointestinal, Hepatobiliary and Transplant Surgery, RWTH Aachen University Hospital, Aachen Germany; NUTRIM School of Nutrition and Translational Research in Metabolism, Maastricht University, Maastricht, the Netherlands; Institute of Pathology, RWTH Aachen University, Aachen, Germany; Department of Surgery, Maastricht University Medical Center, Maastricht, The Netherlands; Department of Medicine III, University Hospital RWTH Aachen, Aachen, Germany; Department of Oncology, University of Oxford, Oxford, United Kingdom; Kennedy Institute of Rheumatology, University of Oxford, Oxford, United Kingdom; Department of Medical Statistics, RWTH Aachen University, Aachen Germany; Department of Pathology, Maastricht University Medical Center+, Maastricht, The Netherlands; CARIM Cardiovascular Research Institute Maastricht, Maastricht University, Maastricht, the Netherlands; Center of Biostatistics and Epidemiology, Medical School, Sigmund Freud University, Vienna, Austria; Department of Hematology, Oncology and Tumor Immunology, CVK, Charité Universitätsmedizin Berlin, Berlin, Germany

**Author notes:** Georg J. Wiltberger and Ulf P. Neumann contributed equally to this work.

**Keywords:** Tumor Microenvironment, Machine Learning, Nerve Fiber Density, Spatial Arrangements, Tertiary Lymphoid Structures

## Abstract

**Background:** B cells and tertiary lymphoid structures (TLS) are reported to be important in the improvement of survival of cancer patients. These secondary lymphoid organs have been associated with the generation of an anti-tumor response. Pancreatic ductal adenocarcinoma (PDAC) is one of the most lethal cancer types and the stromal architecture shapes the intratumoral heterogeneity. The stroma of PDAC is a complex system in which crosstalk takes place between cancer-associated fibroblasts, immune cells, endothelial cells and the cancer cells. Besides immune cells and fibroblasts, there is some limited data about the influence of nerve fibers on cancer progression.

**Patients and methods:** Nerve Fiber Density (NFD) was analysed in our cohort of 188 patients with Pancreatic Ductal Adenocarcinoma who underwent pancreatic surgery. We used immunohistochemistry and multiplex imaging to phenotype the immune cell infiltrate. The cell detection classifier measured distance from immune cell to cancer gland and with a heat map we could count TLS. By using Machine learning we were able to define the spatial distribution and counting Tertiary Lymphoid Structures.

**Results:** High NFD is significantly associated with prolonged overall survival (HR 1.676 (95%CI 1.126,2.495) for low vs. high NFD, p-value 0.0109). The immune cells surrounding the nerve fibers were phenotyped in B cells, T cells and dendritic follicular cells, matching a TLS. Here we show that small nerve fibers are located at the TLS in Pancreatic Cancer and a high Nerve Fiber Density combined with more than 5 TLS is associated with a better survival (HR 0.388 (95%CI 0.218, 0.689).

**Conclusion:** The co-localization of small nerve fibers with TLS is a new finding which has not been described before. However the precise roles of these TLS and nerve fibers remains unknown. These findings unravel future pathways and has the potential to reach new directions into already existing targeted therapy.

## INTRODUCTION

In order to understand the aggressiveness of Pancreatic Ductal Adenocarcinoma (PDAC) and its resistance to therapeutics the components within the cancer stroma needs to be further unravelled. PDAC arises in a tumor microenvironment (TME) that is characterized by extensive communication between tumor cells and non-malignant cells. Specifically, stromal components have been mechanistically implied in immune evasion in the context of cancer therapy, including immunotherapy(*1, 2*). The role of nerve fibers within the cancer-associated stroma has not really been investigated. The aim of this study was to define the role of small nerve fibers in the cancer stroma and their spatial arrangement to immune cells. We hypothesized that small nerve fibers are one of the key components in cancer progression.

Recent publications show that B cells play an important role in survival of cancer patients and response to immunotherapy(*3-7*) and this is not limited to T cells only(*8-10*). Furthermore, B cells play an important role in the tumor formation of PDAC(*11*). Tertiary lymphoid structures have been recognized as ectopic lymphoid organs that reside in inflamed tissue and also in cancer(*12, 13*). These structures show differences in maturation stage and sometimes results in formation of a germinal centre(*13-15*). Their presence is described in multiple cancer types and is variably present in cancer types and patients and is a favourable prognostic factor(*16, 17*).

## RESULTS

### High Nerve Fiber Density is associated with a better survival in Pancreatic Cancer

To gain deeper insight of the influence of nerve fibers on survival we used the neuronal immunohistochemistry staining PGP9.5 on our cohort of patients with pancreatic cancer (n=166). A multivariate Cox regression model adjusted for age, gender and BMI, revealed high Nerve Fiber Density (NFD) (more than 10 positive nerve fibers) to be significantly associated with overall survival (HR 1.676 (95%CI 1.126,2.495) for low vs. high NFD, p-value 0.0109). (see Supplementary Table S1 and Figure S2). While scoring the slides it was observed that the nerve fibers were surrounded by lymphocytes.

### The lymphocyte predominant immunophenotype mainly in a low cellular tumor

To further understand the nerve fiber related immune cells, each routine H&E slide was scored manually into a lymphocyte predominant, neutrophil predominant or immune cell depleted phenotype(*18*). The scoring was done by a senior pathologist and based on the most dominant immune cell on one tumor slide only. It showed that patients with a more abundant stromal phenotype, so a low cellular tumor, have a lymphocyte predominant phenotype. High tumor cellularity was significantly associated with poor survival (HR 4.287 (95%CI 1.460,12.589), p-value 0.0081 for one unit increase). All results are summarized in Table S1 in the Supplementary.

### Immune cells located at the nerve fibers are B cells

Single immunohistochemistry was used to define which immune cells are co-localized with the nerve fibers. The immune cells that were present near the nerve fibers are mainly CD20 and CD8 positive. It is known that PDAC patients with higher levels of CD4+ and CD8+ T cells have a better survival(*19*). TLS are composed of B cells and CD8 positive T cells. To show the maturity of the TLS we used CD21 to define follicular dendritic cells in a germinal centre. In our cohort we found clusters of lymphocytes with and without follicular dendritic cells. This supports the theory of different maturation stages of the TLS(*14*) and mature TLS show the presence of germinal centres. Multiplex immunofluorescence shows the same architecture of a TLS in Pancreatic Cancer (see Figure 1).

**Figure 1:**
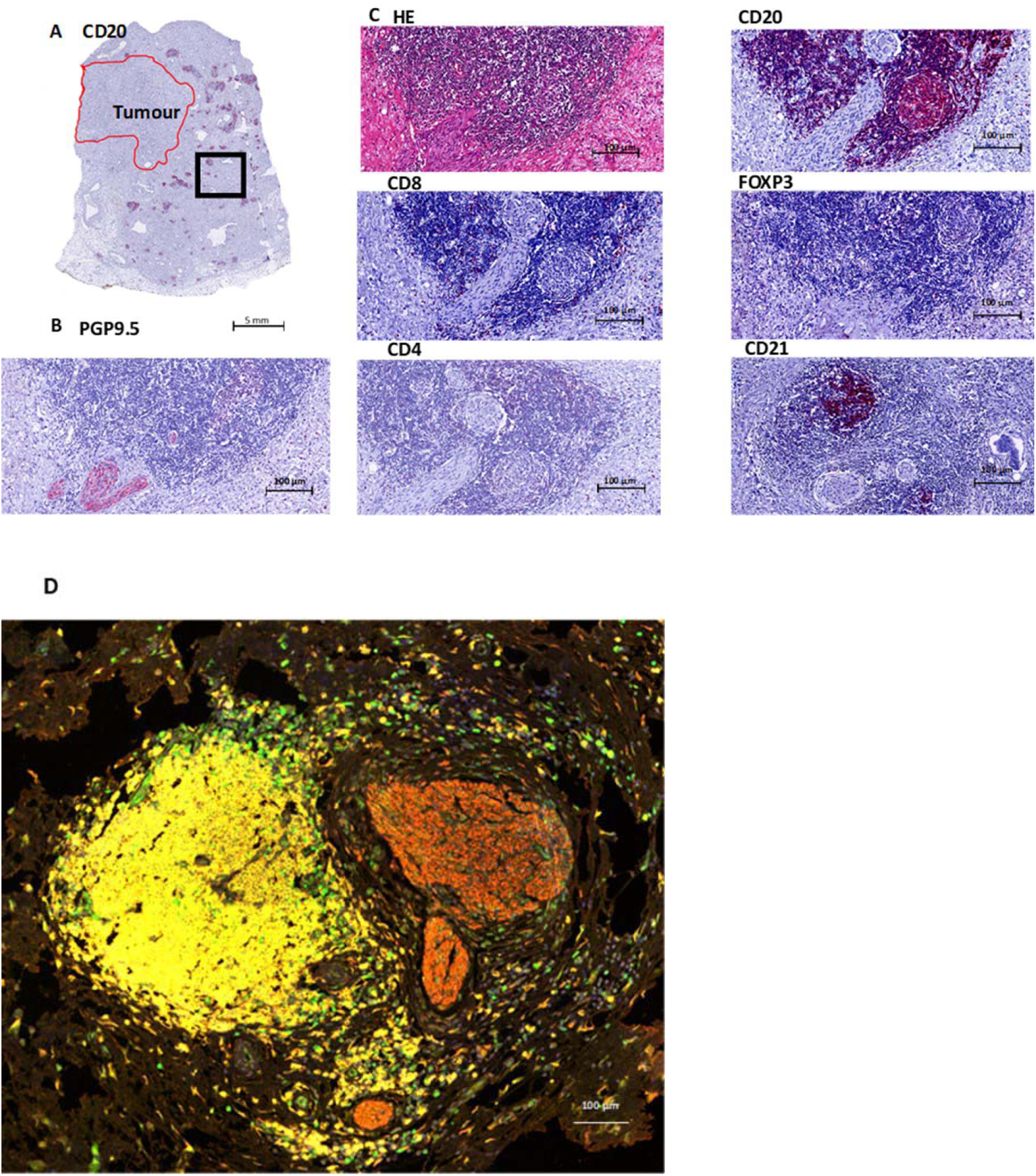
Overview of a TLS in Pancreatic Cancer. **A**. Overview of a CD20 staining(B cells) in a PDAC patient, The red area indicates the tumor region and the black box indicates the area of magnification. **B**.Nerve fiber staining PGP 9.5. **C**.TLS on HE, containing B cells (CD20), T cells (CD8, CD4), and follicular dendritic cells (CD21). Treg cells (FOXP3) are mostly absent, just as CD4. CD8 shows a few positive cells at the border of the TLS. **D**.Multipleximaging: T cells (FOXP3 green), B cells (CD20 yellow), Nerves (PGP9.5 red) and nucleus (DAPI blue).

### Machine learning for quantification of TLS

To gain insight in the spatial arrangements of the immune cells we used machine learning to further measure the distance from the immune cells to the tumor glands. The slides were scanned first and annotations were made in QuPath 0.1.6. A cell detection classifier was trained to recognize immune cells within a region of interest and separate the immune cells from fibroblasts (See Figure 2). To obtain estimates for immune cell densities a kernel-density was applied. The results are displayed as a heat map together with the slides tumor annotations (see Figure 3). To examine the predictive power of the number of tertiary lymphoid structures (despite the non-significant univariate effect) in combination with nerve fibre density and tumor cellularity on overall survival, an additional exploratory Cox regression model with TLS, NFD and TC as well as the interaction between TLS and NFD or TC respectively was evaluated. As the interaction between TLS and TC was non-significant at a 5% level, the interaction was removed from the model. Thus, the exploratory model contains TLS, NFD, TC and the interaction between TLS and NFD as explanatory factors. The significant NFD*TLS interaction (p-value 0.0220) suggests that the effect of NFD is different by TLS. For TLS number greater or equal to 5 mortality is significantly lower in patients with high NFD compared to patients with low NFD (20% (n=14) vs. 10% (n=7); HR 0.388 (95%CI 0.218, 0.689)). Whereas for TLS number less than 5 no significant difference between patients with high or low NFD could be shown (14% (n=13) vs. 19% (n=18); HR 0.959 (95%CI 0.573, 1.604)) which is also apparent in the corresponding Kaplan-Meier plot (See Figure 3).

**Figure 2:**
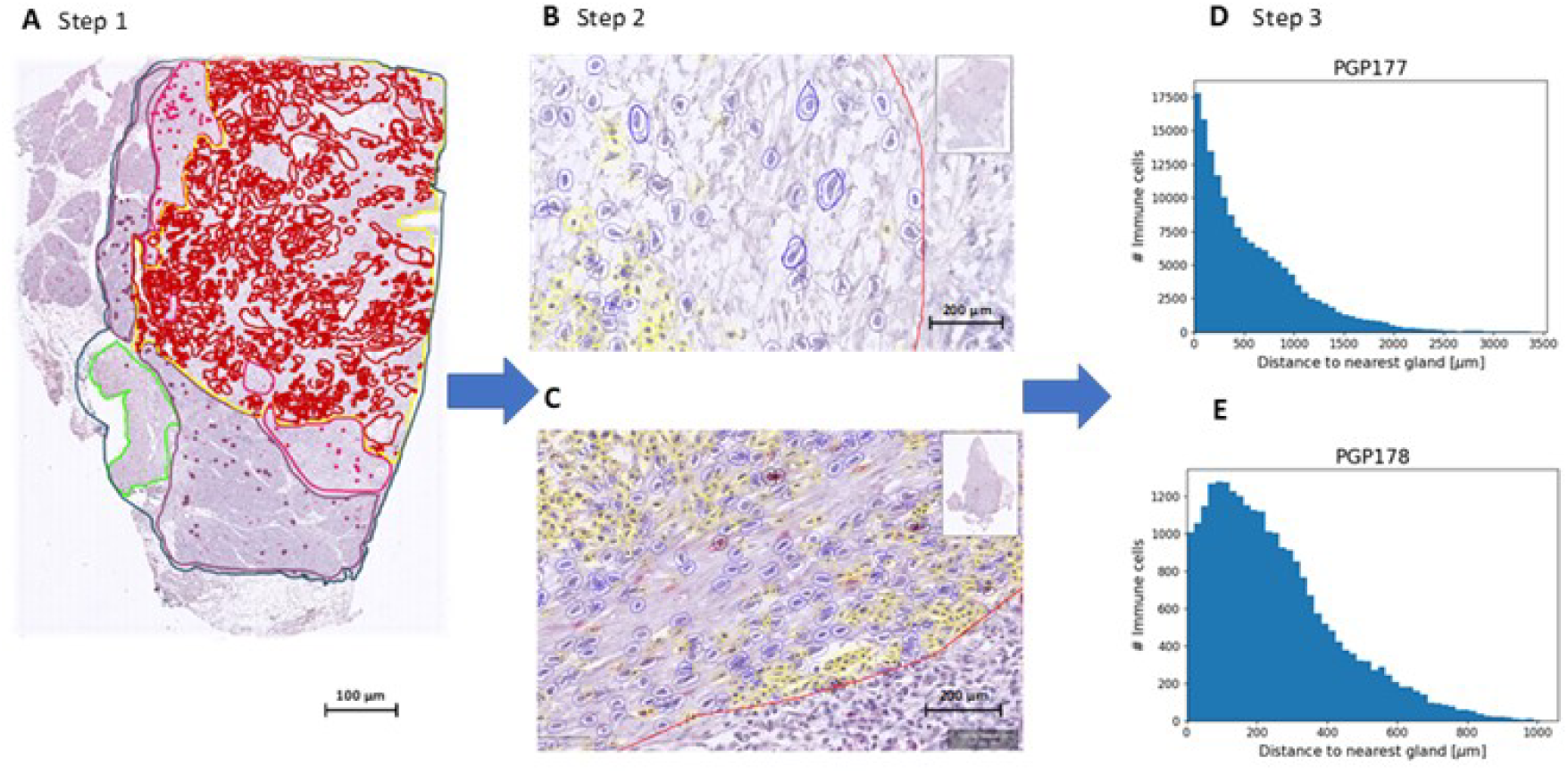
Process using Machine Learning to determine Immune Cell spatiality. **A.** Step 1. Defining the ROI (yellow line). Every scanned slide was annotated in: total tissue (dark blue), normal pancreas (purple), atrophic pancreas (pink), normal duodenum (green) and tumor (red). For the machine learning classifier a Region of Interest (ROI) was also annotated (yellow). In this area only the cell detection classifier was used to detect cells. **B+C.** Step 2. The classifier was trained by the pathologist to recognize fibroblasts (blue) and immune cells (yellow). **D+E.** Step 3. Measurement of distance of Immune Cell to Tumor gland. From each slide a plot was made with amount of immune cells (y-axis) and the distance to nearest tumor gland (x-axis). These plots were used to measure the mean distance from immune cell to tumor gland in micrometer.

**Figure 3:**
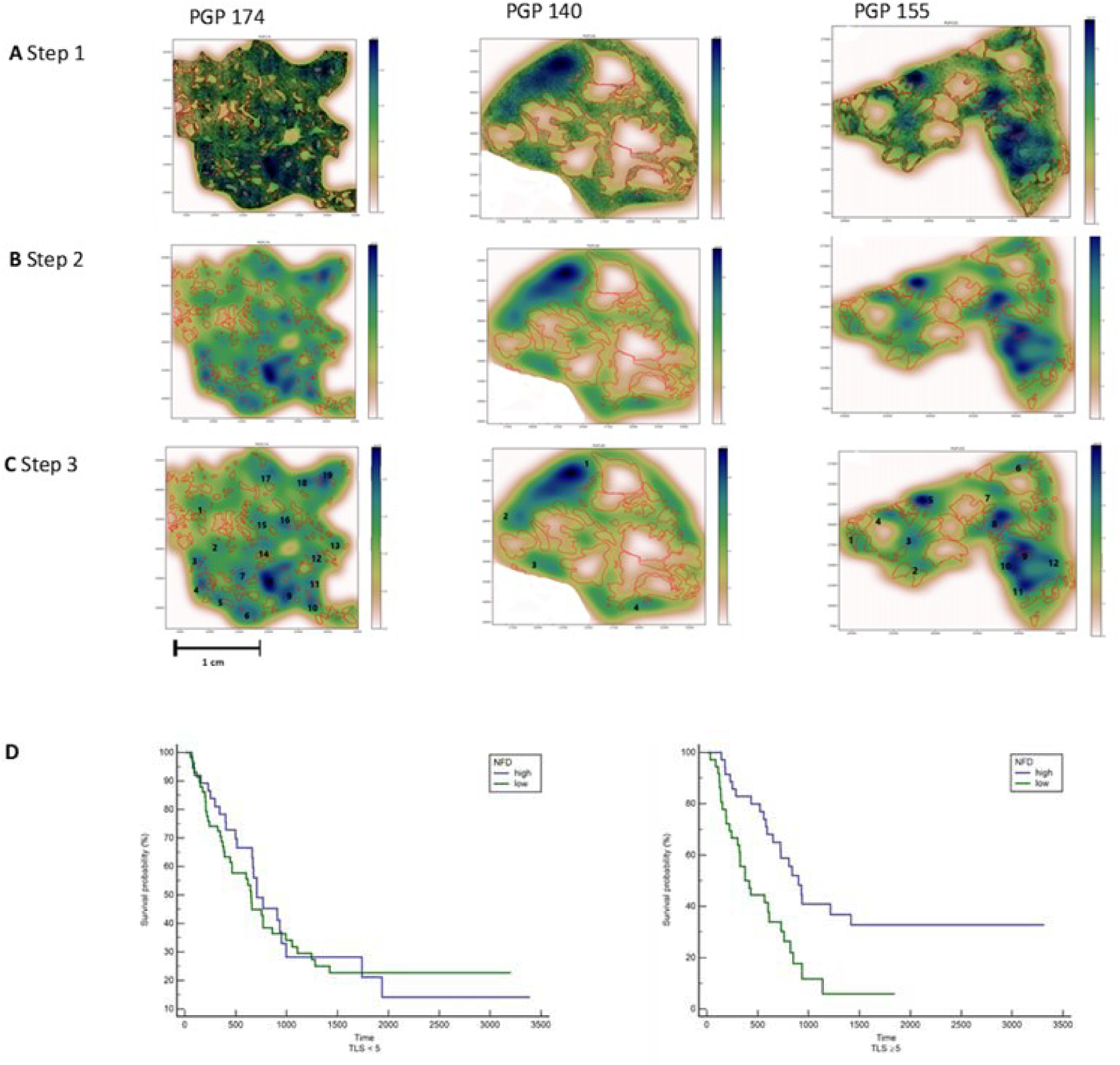
Counting the number of TLS in patients with Pancreatic Cancer. **A.** Step 1 all immune cells are plotted by using the x- and y coordinates. Tumor glands annotated in red. **B.** Step 2 A heat map is created by using a 2D Kernel Density. **C.** Step 3 The heat map clusters with a dark blue and light blue colour are interpreted as a TLS and counted manually. **D.** Left: Kaplan-Meier plot for patients with less than 5 TLS show no significance in survival. Right: Kaplan-Meier plot for patients with 5 or more TLS and a high NFD show a significantly better survival.

## Discussion

PDAC is known for its significant cancer associated stroma or tumor microenvironment. The large stromal component is a significant area of investigation and it is held responsible for poor treatment response. In this study we show that nerve fibers also play a role in the TME. Nerves are emerging regulators of cancer initiation, progression, and metastasis(*20*). Previous data described by Renz et al. suggest that cholinergic signalling by the parasympathetic nerves can suppress the growth of pancreatic cancer cells, where the sympathetic nerves stimulate the growth of pancreatic cells. Therefore, in pancreatic cancer cells there is a balance of neural influence(*21*). Immune cells also play a role in nerves in cancer and are a potential target. There are many levels of neuroimmune interactions, including regulation of inflammation, that play a role in cancer growth and dissemination(*22*).

Neural invasion by tumor cells is one of the most striking characteristics of PDAC and is a sign of aggressive behaviour. Surprisingly in this study we found that a high NFD is associated with a better survival in patients with PDAC. NFD is determined as the amount of small nerve fibers not to be confused by nerves invaded by tumor cells. We found that these small nerve fibers are located in the TLS and predict a better survival in patients with 5 or more TLS and a high NFD.

PDAC is known for its tumor microenvironment and here cross talk takes place between tumor and host. The immune environment is influenced by the tumor and the host. The immune system is known to have a crucial role in cancer and this is possibly regulated by genetic and morphological features of the tumor. Surrounding stromal cells support tumor budding of the cancer cells and promote aggressive behaviour, it is described that this phenotype contains a depletion of TILs (*23*). The presence of tumor infiltrating lymphocytes (TILs) is a predictor of a better prognosis and in breast and ovarian cancer a major component of TILs is the tumor-infiltrating lymphocytic B cells (TIL-B) (*24*). Previous literature has shown that B cells and tertiary lymphoid structures promote immunotherapy response in melanoma and sarcoma (*3-5*) and are of main importance for a better survival. The role of nerve fibers in the co-localization with these TLS has the potential to discover new pathways for a better survival, it could predict sensitivity to immunotherapy and paves the way for new possible targets for (combination)therapy. We expect that these data may be broadly applicable to other malignancies.

## Methods

Histological slides from 166 patients with PDAC were selected and a representative tumor block was retrieved from the archives of the Pathology Department of RWTH University Hospital Aachen, Germany and used to cut tissue sections for the immunostaining Protein Gene Product 9.5 (PGP9.5).

### Patient Cohort

Of the 188 patients included in the cohort 15 were excluded due to in hospital-mortality, 3 due to a loss to follow up, and 4 were excluded due to poor quality pathology slides. Analysis was performed on 166 cases. Clinical data for this cohort are listed in Supplementary Information Table S1.

### Pathological Examination

The clinic-pathological parameters, including tumor size, differentiation, positive lymph node status, R0/R1, were carefully reviewed in the original report. All PDAC lesions were pathologically examined and classified according to World Health Organization (WHO) classification using the TNM Classification for Malignant Tumors.

#### Nerve fiber density (NFD)

Immunohistochemistry was performed on formalin-fixed, paraffin embedded tissue sections as described previously. Sections (2.5μm thick) were cut, deparaffinized in xylene and rehydrated in graded alcohols. Slides were boiled in citrate buffer (pH 6.0) at 95 - 100°C for 5 minutes and were cooled for 20 minutes. Endogenous peroxide in methanol for 10 minutes. Sections were incubated with rabbit anti-human PGP 9.5 (DAKO 1:100) overnight at 4°C.

A single digital image was uploaded in Qupath 0.1.6 which is a flexible software platform suitable for a range of digital pathology applications. As previously described all slides were read by a senior pathologist and NFD was evaluated by counting the number of nerve fascicles with diameters of <100 μm in 20 continuous fields at x 200 magnification.

Nerve fiber density results were grouped into 3 categories: 1) negative, no nerve fibres, 2) weak expression, 1-10 nerve fibres and 3) moderate/strong expression >10 nerve fibers, according to existing literature on breast cancer(*25*).

#### Tumor Cellularity (TC)

Every immunostained slide was scanned and on whole slide imaging the tumor glands, normal pancreatic tissue and atrophic pancreatic tissue were manually annotated in QuPath 0.1.6 by a senior pathologist[18]. Stromal area was measured by following formula: total tissue - (normal tissue + atrophic pancreas+ tumor) = stroma surface. Tumor cellularity was measured as (tumorsurface / (tumorsurface + stromasurface).

#### Phenotyping of Immune Cells

An H&E slide was cut from the paraffin embedded tissue and the pre-dominant type of immune cells was judged manually by the PhD student and the senior pathologist. The H&E slide and the slide used for immunostaining were cut directly after each other. The dominant type of immune cells was determined by the senior pathologist on the corresponding H&E slide and manually scored into three categories: 1) lymphocyte predominant, 2) neutrophil predominant or 3) no immune cells.

#### Single Immunohistochemistry

To further define the type of immune cell, we performed immunohistochemistry on 10 patients with histologically confirmed TLS and a high NFD. We used CD20 (B cells), CD4, CD8, FOXP3 (all T cell markers), according to previous literature that CD20+ B cells were located in the TLS and were colocalized with CD4+, CD8+, and FOXP3+ T cells(*5*). To illustrate the presence of germinal centres we also stained for CD21[19]. The stained slides were compared to the H&E routine staining and PGP9.5 nerve fiber staining (examples see Figure 1).

H&E and immunohistochemistry staining were performed on FFPE tumor tissue sections. The tumor tissues were fixed in 10% formalin, embedded in paraffin, and serially sectioned.

Sections were stained with mouse or rabbit anti-human monoclonal antibodies against CD20 (Dako, L26, 1:200), CD21 (DAKO, 1:25, CD23 (Leica, CD23-1B12, 1:50), CD4 (DAKO, 4B12 1:50), CD8 (DAKO, C8/144B, 1:50), FOXP3 (DAKO, PCH101, 1:50). All sections were counterstained with haematoxylin, dehydrated and mounted. All sections were cover slipped using Vectashield Hardset 1500 mounting medium with DAPI and slides were scanned and digitalized using the Roche Ventana scanner. Immunohistochemistry staining was interpreted in conjunction with H&E stained sections.

#### Multiplex immunofluorescence assay and analysis

For images shown in Fig. 1, immunofluorescence multiplex staining, we followed the staining method for the following markers: CD20 (Dako, L26, 1:500) with subsequent visualization using fluorescein Cy3 (1:100); FOXP3 (DAKO, PCH101, 1:300) with subsequent visualization using fluorescein FITC (1:100); PGP9.5 (DAKO 1:300) with subsequent visualization using fluorescein TEX RED (1:100) and nuclei visualized with DAPI. All of the sections were cover slipped using Vectashield Hardset 1500 mounting medium.

The slides were scanned using the TissueFAXS slide scanner (supplier TissueGnostics). For each marker, the mean fluorescent intensity per case was then determined as a base point from which positive cells could be established. For multispectral analysis, each of the individually stained sections was used to establish the spectral library of the fluorophores. The senior pathologist selected the Region of Interest (ROI) at 20× magnification.

#### TLS count

By using the annotations on the immunostained slide (PGP9.5), the distance from each immune cell to tumor gland was measured. The spatial arrangement of the immune cells was determined by a semi-automated machine learning workflow, which comprised cell segmentation, feature computation and stroma- and immune cell identification. To facilitate high trough put of thousands of immune cells on multiple images, QuPath enables interactive training of cell classification, after which the classifier can be saved and run over multiple slides. The senior pathologist trained a cell detection classifier to recognize immune cells and fibroblasts in a certain Region of Interest (ROI). This ROI was annotated by the senior pathologist and contained tumor glands and stroma only, it was avoided to include normal tissue and/or atrophic tissue in order to achieve the best detection results. Application of this workflow resulted in both fine grained cell-by-cell analysis and overall summary scores of the spatial arrangements of the immune cells within the ROI in relation to the annotated tumor glands. The results are visualized via color-coded markup images. Representative examples of these markup images are shown in Figure 2.

To obtain estimates for immune cell densities a kernel-density estimate using Gaussian kernels as implemented in the gaussian_kde class from SciPy is applied. First kernels are fitted from immune cells positions and calculated for points on a 100×100 grid for each slide. Subsequently the results are displayed as a heat map together with the slides tumor annotations (see Figure 3). From this heat map regions of high immune cell density can be identified manually that correspond to a TLS.

#### Statistics

Continuous variables were summarized by means and corresponding standard deviations (SD). Categorical data were presented by frequencies and percentages. Cox regression models were used to analyse the joint relation between clinical variables (coded by 0 and 1 for binary variables) on overall mortality. All exploratory variables were studied in an univariate Cox regression model adjusted for age, gender and BMI. Exploratory variables were assessed as relevant to be mutually included in our final model if the p-value was below 0.05. Relevant variables were studied further for pairwise interaction. In doing so, we used again the significance margin of 5%. Then a multivariate Cox regression model adjusted for age, gender and BMI with backward selection was fitted to the previously identified variables and interactions. During this final step, the significance level for removing a variable or interaction from the model was set to 0.05. For the final Cox model, graphical and numerical methods according to Lin et al. were performed to establish the validity of the proportionality assumption (*26*). No deviation from model assumption could be observed. We report our results by hazard ratios, corresponding 95% confidence limits and p-values, where a p-value of less or equal than 0.05 could be interpreted as statistically significant test results. Forest plots were chosen for graphical visualization and Kaplan-Meier plots for comparison of subgroups. All analyses were performed using SAS^®^statistical software, V9.4 (SAS Institute, Cary, NC, USA).

## Ethics statement

All experiments were conducted in accordance with the Declaration of Helsinki and the International Ethical Guidelines for Biomedical Research Involving Human Subjects. Anonymized archival tissue samples were retrieved from the pathology archive at the University Hospital of Aachen (RWTH Aachen) after approval by the institutional ethics board under protocol number EK 106/18.

## Data availability

Extended data is available for this paper at https: (summary tables with raw data will be made available after acceptance.)

Supplementary information is available for this paper and link will follow after acceptance.

## Code availability

Source codes are available at https://github.com/janniehues/PGP-pancreas/

## Authors’ Contributions

**Conception and Design:** L.R. Heij, U.P. Neumann

**Data analysis and interpretation:** L.R. Heij, X. Tan, G.J. Wiltberger, J.N. Kather, J.M. Niehues, G. van der Kroft, M. Aberle

**Annotation of slides:** X. Tan and L.R. Heij

**Design and implementation of the algorithm:** J.N. Kather, J.M. Niehues and J. Cleutjens

**Statistical analysis:** G.J. Wiltberger and N. Heussen

**Writing, review and/or revision of the manuscript:** L.R. Heij, G.J. Wiltberger, J. Bednarsch, U.P. Neumann, S. Sivakumar, D.P. Modest

**Material support and histology expertise:** N. Gaisa and D. Liu

**Study supervision:** T. Luedde, S. Olde Damink, U.P. Neumann and S. Lang

All authors reviewed and approved the final manuscript as submitted and agree to be accountable for all aspects of the work.

The authors declare no competing interests.

### Online content

Any methods, additional references, source data, extended data, supplementary information, acknowledgements, peer review information, details of author contributions and competing interests; and code availability is available at https://github.com/janniehues/PGP-pancreas/

The authors declare no potential conflicts of interest

