## Supplementary Information for "Nerve fibers in the Tumor Microenvironment are co-localized with Tertiary Lymphoid Structures"

|  |  | Univariate analyses  (adjusted for age, gender, BMI) | | Multivariate analysis  (adjusted for age, gender, BMI) | |
| --- | --- | --- | --- | --- | --- |
| Variable | Mean (SD) or  Frequency (%) | Hazard ratio [95%-C]) | p-value | Hazard ratio (95%-CI) | p-value |
| Age | 66 (10) |  |  | 1.024  [1.002; 1.047] | 0.0356 |
| Gender  female | 80 (48.19) |  |  | 1.173  [0.789; 1.745] | 0.4297 |
| male | 86 (51.81) |  |  |  |  |
| BMI | 25.7 (4.3) |  |  | 0.996  [0.954; 1.040] | 0.8547 |
| ASA  $<3$ | 62 (37.35) | 0.909  [0.607, 1.362] | 0.6449 |  |  |
| $\geq3$ | 104 (62.65) |  |  |  |  |
| Tumor grade  G2 | 94 (56.63) | 1.954  [1.342, 2.844] | 0.0005 |  |  |
| G3 | 72 (56.63) |  |  |  |  |
| Extent of tumor  T1/T2 | 26 (15.66) | 2.116  [1.015, 4.410] | 0.0455 |  |  |
| T3/T4 | 140 (84.34) |  |  |  |  |
| Perineural invasion  Absent | 28 (16.87) | 2.239  [1.265, 3.961] | 0.0056 | 2.409  [1.337; 4.340] | 0.0034 |
| Present | 138 (83.13) |  |  |  |  |
| Lymph node metastasis  Absent | 39 (23.49) | 2.322  [1.407, 3.834] | 0.0010 |  |  |
| Present | 127 (76.51) |  |  |  |  |
| Lymphatic invasion  Absent | 114 (68.67) | 2.080  [0.418, 3.050] | 0.0002 | 1.763  [1.173; 2.651] | 0.0064 |
| Present | 52 (31.33) |  |  |  |  |
| Venous invasion  Absent | 136 (81.93) | 1.452  [0.923, 2.284] | 0.1068 |  |  |
| Present | 30 (81.93) |  |  |  |  |
| Surgical margin status  Negative | 106 (63.86) | 1.962  [1.342, 2.868] | 0.0005 |  |  |
| Positive | 60 (36.14) |  |  |  |  |
| Nerve fibre density  High | 72 (43.37) | 1.597  [1.093, 2.336] | 0.0155 | 1.676  [1.126, 2.495] | 0.0109 |
| Low | 94 (56.63) |  |  |  |  |
| TLS  $<5$ | 95 (57.23) | 1.084  [0.745, 1.577] | 0.6723 |  |  |
| $\geq5$ | 71 (42.77) |  |  |  |  |
| Tumor cellularity | 0.36 (0.20) | 5.280  [1.952, 14.282] | 0.0010 | 4.287  [1.460; 12.589] | 0.0081 |
| Interaction between Lymph node metastasis and surgical margin status |  |  |  |  | 0.0053 |
| lymph node metastasis present at surgical margin status positive |  |  |  | 0.587  [0.272; 1.266] |  |
| lymph node metastasis present at surgical margin status negative |  |  |  | 2.618  [1.260; 5.437] |  |

*Table S1: Results from univariate and multivariate Cox regression analysis adjusted for age, gender and BMI of prognostic factors associated with overall survival*

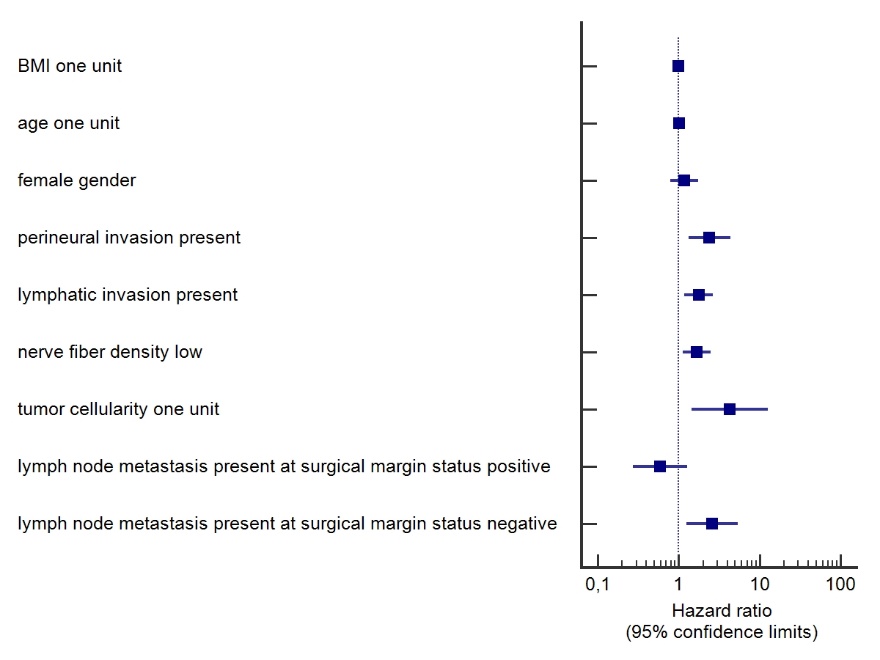

Figure S1: Forest plot of hazard ratios and corresponding 95% confidence limits

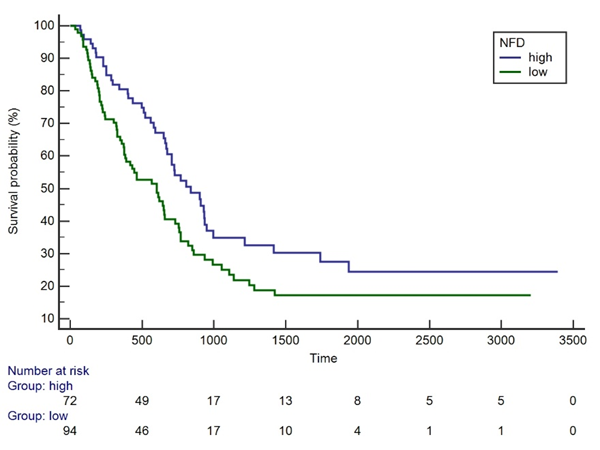

Figure S2: Kaplan-Meier plot of the survival probabilities of patients with high and low NFD
